# An ATP-Binding Cassette Transporter Gene Links Innate and Adaptive Immune Responses

**DOI:** 10.1101/2023.12.31.573785

**Authors:** Sara M. Wilcox, Hitesh Arora, Kyung Bok Choi, Suresh Kari, Lonna Munro, Cheryl G. Pfeifer, Wilfred A. Jefferies

## Abstract

Positive-strand RNA viruses and DNA viruses generate double-stranded RNA (dsRNA) during their replication processes and innate immune responses against viral infections are orchestrated by numerous interferon-stimulating genes, yet the detailed coordination of downstream signaling of anti-viral immune responses is not fully understood. Recent studies suggest 2’-5’- Oligoadenylate Synthetase 1 (OAS1) may have a protective role in severe acute respiratory syndrome coronavirus 2 (SARS-CoV-2) infections; however, the mechanism regulating OAS1 remains uninvestigated. Our aim is to understand the regulation of OAS1 and its modulation of RNaseL activity, as this has significant implications for responses to RNA viruses, including Vesicular stomatitis virus (VSV) and SARS-CoV-2. We explore the hypothesis that ABCF1 an ATP-binding cassette family member protein, a key regulator of innate immune responses and macrophage polarization and cytokine storm, play a role in regulating the antiviral responses and downstream dsRNA signaling revealed by measuring responses to the synthetic dsRNA analog termed poly (I:C). We utilize ABCF1 haplo-insufficient mice to discover that ABCF1 modulates the amplitude and frequency of VSV-specific Cytolytic T lymphocyte in anti-viral immune responses and suggests that innate immune responses underpin this process. To understand this mechanism, we describe that ABCF1 interacts with 2’-5’-oligoadenylate synthetase 1 (OAS1) which in turn modulates essential proteins that leads to the modulation of RNaseL activity via ABCE1. Furthermore, we find that ABCF1 also influences the production of interferon-α (IFN-α) and interferon-β (IFN-β) in bone marrow-derived macrophages. Overall,, we unexpectedly discovered that ABCF1 acts as a crucial link between innate and adaptive immunity, regulating the development of adaptive Cytolytic T lymphocyte responses and interacting with OAS1, a key regulator of innate immune responses against viral infections. Exploring pharmacological agents that target ABCE1 or ABCF1 may lead to the discovery of novel modalities for countering SARS CoV-2 and other viruses where OAS1 is a crucial innate immune response gene.

## Introduction

The innate immune system drives adaptive immunity and is the critical first line of defense against viral infections including SARS CoV-2 (1–3). It does so by sensing the infection through pattern recognition receptors (PRR). The endosomal Toll-like receptors (TLRs 3,7/8) recognize double stranded RNA (dsRNA) and single stranded RNA (ssRNA) respectively. Synthetic dsRNA analog polyriboinosinic-polyribocytidylic acid (poly(I:C)) is a potent ligand of TLR3 and that can also activate RIG-I, MDA-5 and PKR (2, 4, 5). Once activated, TLR3 recruits TIR-domain-containing adapter-inducing interferon-β (TRIF) (Anti TRIF antibody sourced from abcam; catalog ab13810). Subsequently, it binds tumor necrosis factor receptor-associated factor 3 (TRAF3) (Anti TRAF3 antibody sourced from Santa Cruz Biotech; catalog sc-6933). This binding activates TRAF3, which further leads to phosphorylation and activation of TANK binding kinase1 (TBK1) (Phospho-TBK1 antibody sourced from abcam; catalog ab109272) (Anti TBK1 antibody sourced from abcam; catalog ab40676) and Inhibitory kappa kinase (IKKe). This results in phosphorylation of transcription factor Interferon regulatory factor 3 (IRF3). IRF3 then translocates in the nucleus, dimerizes and leads to transcription of type I interferons (IFN-I) (6).

The binding of IFN I receptor leads to rapid auto-phosphorylation of receptor associated tyrosine Kinase 2 (TYK2) and janus activated kinase 1(JAK1). This activation phosphorylates and activates signal transducer and activator of transcription (STAT proteins) (7, 8). The major STATs that are induced by IFN-I are STAT1 and STAT2 that form a complex with IRF9, which is termed IFN-stimulated genes (ISG)-F3 (9, 10). These STAT proteins form either homodimers or heterodimers and these complexes bind to IFN-stimulated response element (ISRE) in IFN stimulating genes (ISG) promoters, thereby initiating their transcription (11, 12). Mitogen activated protein kinases (MAPK) are also known to mediate IFN I signaling via phosphorylation of p38 and ERK kinases (13). Studies have shown that MAPK kinases are independent of STAT pathway, as the inhibition of p38 activity does not block STAT or ISGF3 activity (14). Downstream p38 and ERK leads to phosphorylation and activation of MSK and MNK kinases, which lead to phosphorylation of various transcription factors (15, 16). The initiation of viral response and production of anti-viral response and cytokines and interferon’s (IFN) is well studied but the mechanism that regulates it remains vague.

One of the anti-viral mechanisms recently discovered is mediated by family of anti-viral proteins called 2’-5’ oligoadenylate synthetases (OAS). Like other anti-viral genes, transcription of OAS1 is induced by virus infection and IFN stimulation (17). In humans there are three functional OAS genes (OAS1-3), which result in 8 to 10 OAS isoforms due to mRNA splicing. In mice, there are 7 separate OAS1 genes, in addition to OAS2 and OAS3 genes (18, 19). After stimulation with dsRNA, the functional OAS proteins exhibit a characteristic polymerase activity and produce a series of short 5’ phosphorylated, 2’,5’-linked oligoadenylates collectively referred to as 2-5A, from ATP (20). 2-5As subsequently bind and activate Ribonuclease L (RNaseL). Activated RNaseL leads to degradation of both cellular and viral RNA, leading to termination of protein synthesis and the cessation of viral replication (21, 22). Studies have also shown that RNaseL enhances the induction of IFNα, thereby controlling viral infection (23). This process has been shown to be regulated through the binding of small RNA cleavage products with RIG-I (Retinoic Acid Inducible Gene I) and MDA5 (Melanoma Differentiation-Associated Protein 5) (21, 24). After production, IFN-β engages with IFNAR1 and IFNAR2, and this binding initiates a signaling cascade - described above - leading to the induction of more than 300 IFN-stimulated genes (ISG’s) (25).

Furthermore, recent studies have been able to demonstrate that an increased Neanderthal OAS1 isoform level in patients of European ancestry is strongly associated with a reduced risk of very severe COVID-19 infection, hospitalization and susceptibility, inferring on them protective immunity against the virus through ribonuclease L (RNase L) (1, 26). This further highlights the importance of developing drugs that target increased OAS1 levels (2, 3).

Differential display PCR identified up-regulation of a novel ABC gene, named ABCF1 in TNF-stimulated synoviocytes in rheumatoid arthritis patients (27). ABCF1 was the first mammalian ABC transporter to be discovered that lacked a trans-membrane domain and doesn’t function as a transporter. The ABCF1 gene is located in the class I region of the MHC locus on chromosome 17 in mice and on chromosome 6 in humans. ABCF1 is also thought to be the mammalian homolog of the yeast protein GCN20 and is also known to help in eIF2 recruitment to 40S ribosomal subunit and initiate translation (28, 29). Immunological studies have shown that ABCF1 is associated with the susceptibility to rheumatoid arthritis in European and Asian populations and to autoimmune pancreatitis in Japanese population (30). Recent studies in mouse embryonic fibroblasts have shown that ABCF1 associates with dsDNA and DNA sensing components HMGB1 and IFI204, and further interacts with SET complex members (SET, ANP32A and HMGB2) to facilitate cytosolic DNA sensing mechanisms (31). Our recent findings have revealed that ABCF1 possesses E2 ubiquitin enzyme activity, which plays a crucial role in the regulation of the immune response to gram-negative insults mediated by Lipopolysaccharide (LPS) and Toll-like Receptor-4 (TLR4) (32). Specifically, ABCF1 targets key proteins for K63-polyubiquitination, a post-translational modification process. This ubiquitination by ABCF1 leads to a shift in the inflammatory profile triggered by LPS-TLR4 signaling. It transitions the response from an early-phase MyD88-dependent pathway to a late-phase TRIF-dependent signaling pathway. Furthermore, ABCF1’s ubiquitination activity influences TLR4 endocytosis, a cellular process of internalization, and has a notable impact on macrophage polarization, shifting it from an M1 to an M2 phase. This transition in macrophage phenotype is significant in regulating the immune response. In physiological terms, ABCF1 plays a pivotal role in orchestrating the shift from the initial inflammatory phase of sepsis to a state of endotoxin tolerance, ultimately attenuating the cytokine storm associated with this condition (32). Additionally, the ATP binding cassette family E member 1 (ABCE1) is a member of ATP cassette super family and is a known RNaseL inhibitor (33). ABC transporters are a ubiquitous family of transport proteins, which is one of the largest and most ancient families, with representatives in all extant phyla from prokaryotes to humans (34). Studies have shown that overexpression of ABCE1 in HeLa cells inhibits IFN mediated anti-viral activity on encephalomyocarditis, thereby inhibiting 2-5A/RNaseL pathway (35).

Positive-strand RNA viruses such as SARS CoV-2 and VSV and DNA viruses, such as Adenoviruses, Herpes viruses and Poxviruses generate double-stranded RNA (dsRNA) during their replication processes (36). Indeed, double-stranded RNA (dsRNA) molecules are a key hallmark of certain viral infections. Positive-strand RNA viruses, which have a genome that can be directly translated into proteins, often generate dsRNA intermediates during their replication process. Similarly, DNA viruses can produce dsRNA during various stages of their life cycle. In contrast, negative-strand RNA viruses, whose genome cannot be directly translated into proteins, typically do not produce detectable amounts of dsRNA (36).

This disparity in dsRNA production among different classes of viruses is a significant factor in how the immune system recognizes and responds to viral infections. dsRNA is a potent trigger for antiviral responses, particularly through the activation of innate immune pathways mediated by receptors like TLR3 and RIG-I and MDA5. These receptors recognize dsRNA and initiate signaling cascades that lead to the production of interferons and other antiviral proteins, aiding the host in combating the viral invasion. Therefore, the presence or absence of dsRNA serves as a critical factor in the immune responses mounted against different types of viruses (36).

Previous research has demonstrated that dsRNAs originating from SARS-CoV-2, an Alphavirus, or cells susceptible to SARS-CoV-2, are internalized by mononuclear phagocytes. This internalization simultaneously triggers two other innate immune pathways: the RIG-I/MDA5-MAVS pathway, leading to the production of inflammatory cytokines, and the OAS-RNase L pathway (Lee et al. (3)).

Here we study VSV, another Alphavirus as well as poly (I:C) stimulation that simulates a dsRNA response, to address the hypothesis whether ABCE1 and ABCF1 can regulate the anti- viral activity and modulate RNaseL activity and therefore responses to positive-sense single-stranded RNA viruses such as VSV and SARS CoV-2. During these studies we make the unexpected discovery that ABCF1 links innate and adaptive immunity by regulating the generation of Cytolytic T lymphocyte adaptive immune responses and interacts with OAS1, a key regulator of innate immune responses to viruses.

## Results

### CD4 and CD8 staining on the spleen and thymus

*Abcf1*^+/−^ mice (Het; B6.Cg-Abcf1 < Gt(XK097)Byg) mice were generated by our laboratory as previously described (37) and C57BL/6J (Jackson Laboratories; 000664) mice were used as wild-type (WT) controls for *in vivo* experiments. It is important to note that complete loss of ABCF1 is lethal because it also appears to have a function in development (37). Therefore, homozygotic knockout models could not be utilized.

To address whether ABCF1 effected lymphocyte development, CD4 and CD8 staining on the spleen and thymus was undertaken and analysed by flow cytometry (**Figure 1**). The thymocyte subpopulations appear to be unaffected in ABCF1+/- compared to controls however, there is an increase in CD4+ cells in spleens of ABCF1+/- compared to controls and because of their size it suggests that these are resting CD4+ T Lymphocytes.

**Figure 1.**
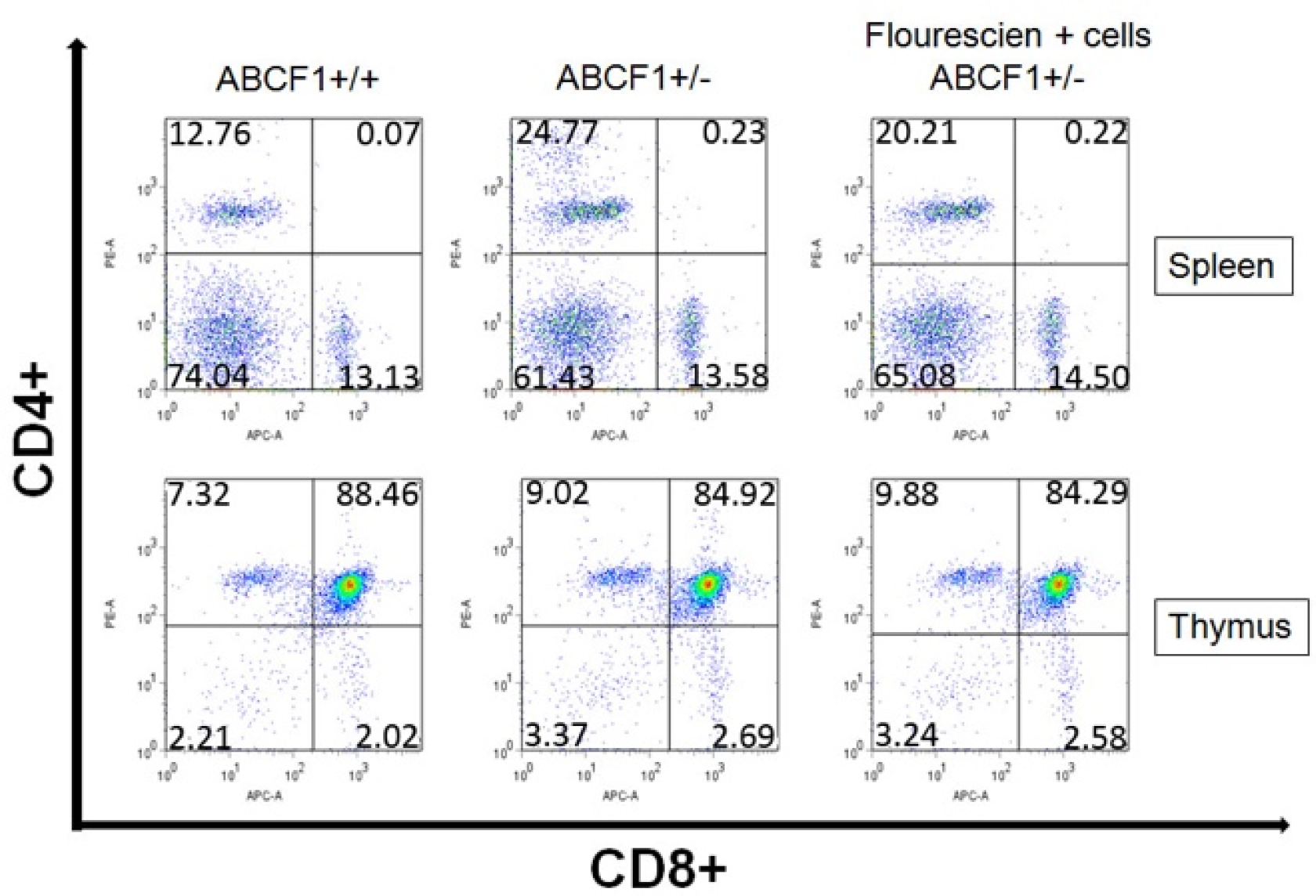
**ABCF1 promoter activity in CD8+ T cells from the thymus and the spleen.** Spleens and thymi were dissected from threeABCF1+/- (top panels) and three of their wild-type littermate controls (bottom panels). Cells were labelled with the αnti-CD8 antibody and αnti-CD4 antibody. We depict the staining of two ABCF1+/- and one wild-type littermate controls. Data is representative of two separate experiments.

### The ABCF1 promoter is active in CD8+ T lymphocytes

Since there was significant background staining in the X-gal-stained tissue sections, we decided to analyse thymocytes using fluorescein di-β-D-galactopyranoside (FDG) staining by FACS to see if we could eliminate the background. Initially we chose to look at the ABCF1 promoter activity in CD8+ T-cells. The thymus is known to be a primary lymphoid organ where bone marrow derived progenitor cells undergo a series of differentiation steps to form mature CD8+ T cells.

In the thymus, approximately 65% of the CD8 positive cells were fluorescein negative and 35% were positive (**Figure 2**). In the spleen in contrast, 84% of the CD8 positive cells were fluorescein positive and only 16% of the cells were fluorescein negative. These data suggest that the ABCF1 promoter could be turned on during the maturation of CD8+ cells (**Figure 2**).

**Figure 2.**
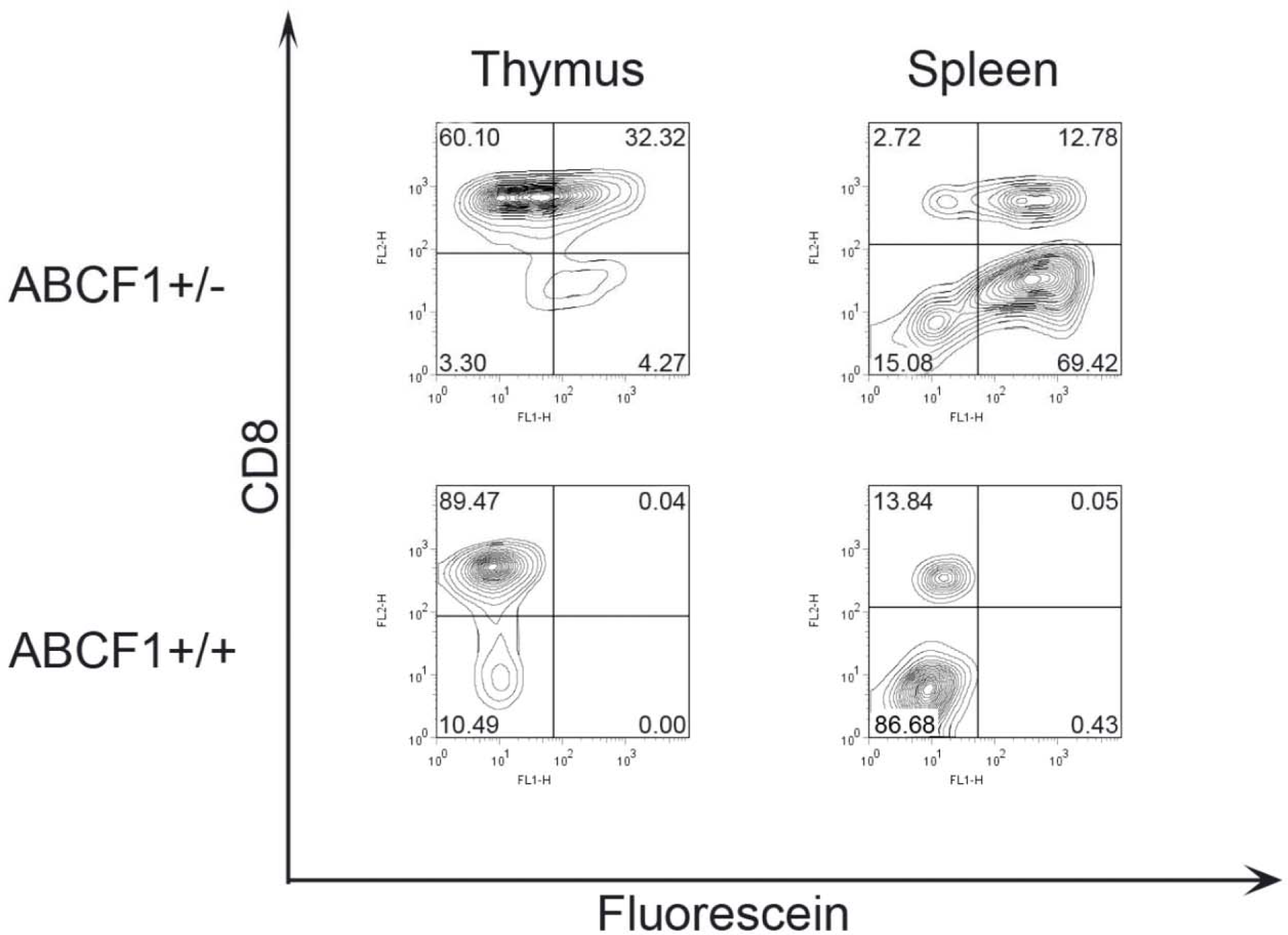
ABCF1 promoter activity in CD8+ T cells from the thymus and the spleen. Spleens and thymi were dissected out of three ABCF1+/- (top panels) and three of their wild-type littermate controls (bottom panels). Cells were labelled with the PE-αCD8 antibody and then stained with fluorescein by heat shock. Data is representative of two separate experiments.

### ABCF1+/- mice produce fewer functional cytotoxic T lymphocytes (CTLs)

To determine whether ABCF1 is important for CD8+ T lymphocyte maturation, a CTL assay was used to assess whether ABCF1+/- mice can produce functional CTLs. After expansion, ABCF1+/- spleens had similar percentages of CD4+ cells but fewer CD8+ T lymphocytes (**Figure 3**). The tetramer and IFNγ staining also revealed that the ABCF1+/- mice had fewer tetramer-specific (**Figure 4**) and fewer IFNγ-producing CD8+ T lymphocytes (**Figure 5**). For the CTL assay, ABCF1+/- and ABCF1+/+ splenocytes were incubated with 51Cr labelled targeT lymphocytes (RMA-S) at various ratios and killing was monitored by the release of 51Cr from the targeT lymphocytes into the supernatant. ABCF1+/- CTLs were found to be as good as ABCF1+/+ CTLs at killing their targets (**Figure 6**).

**Figure 3.**
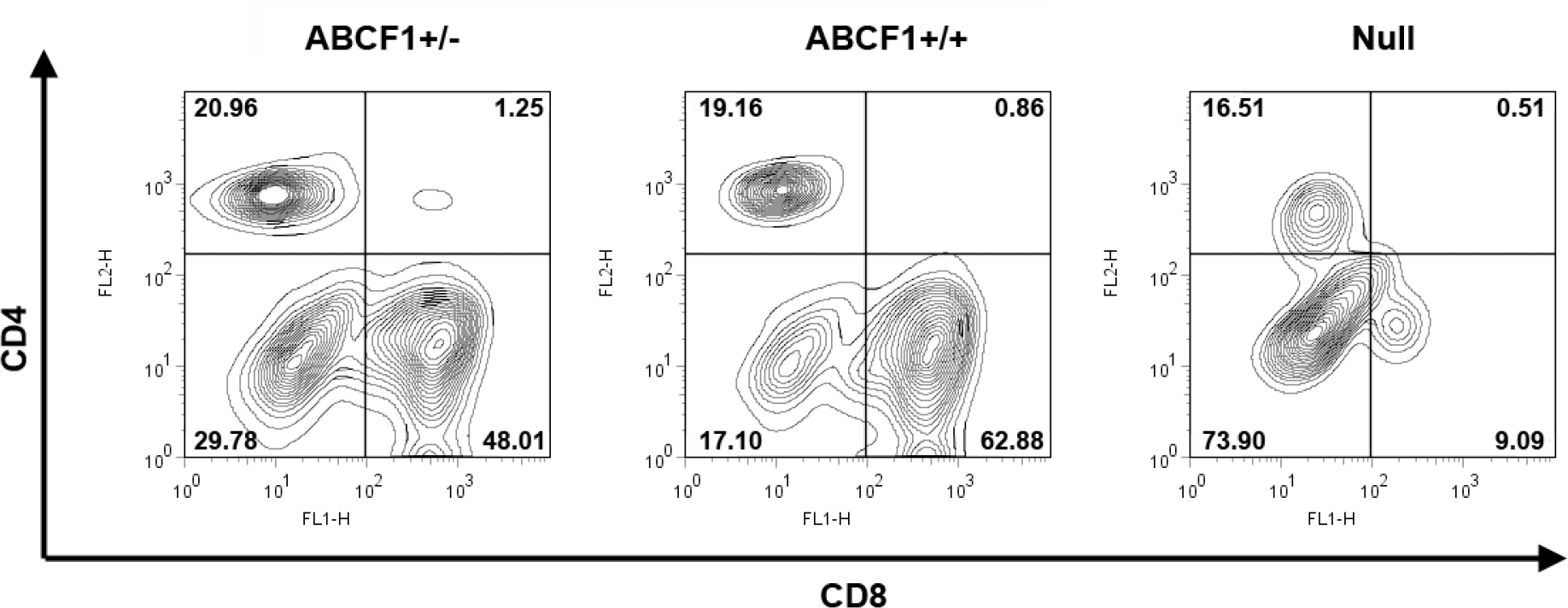
ABCF1 +/- mice produce fewer CD8+ T lymphocytes after infection with VSV. Figure shown represents the results of one experiment, which was later repeated seven times with similar results.

**Figure 4.**
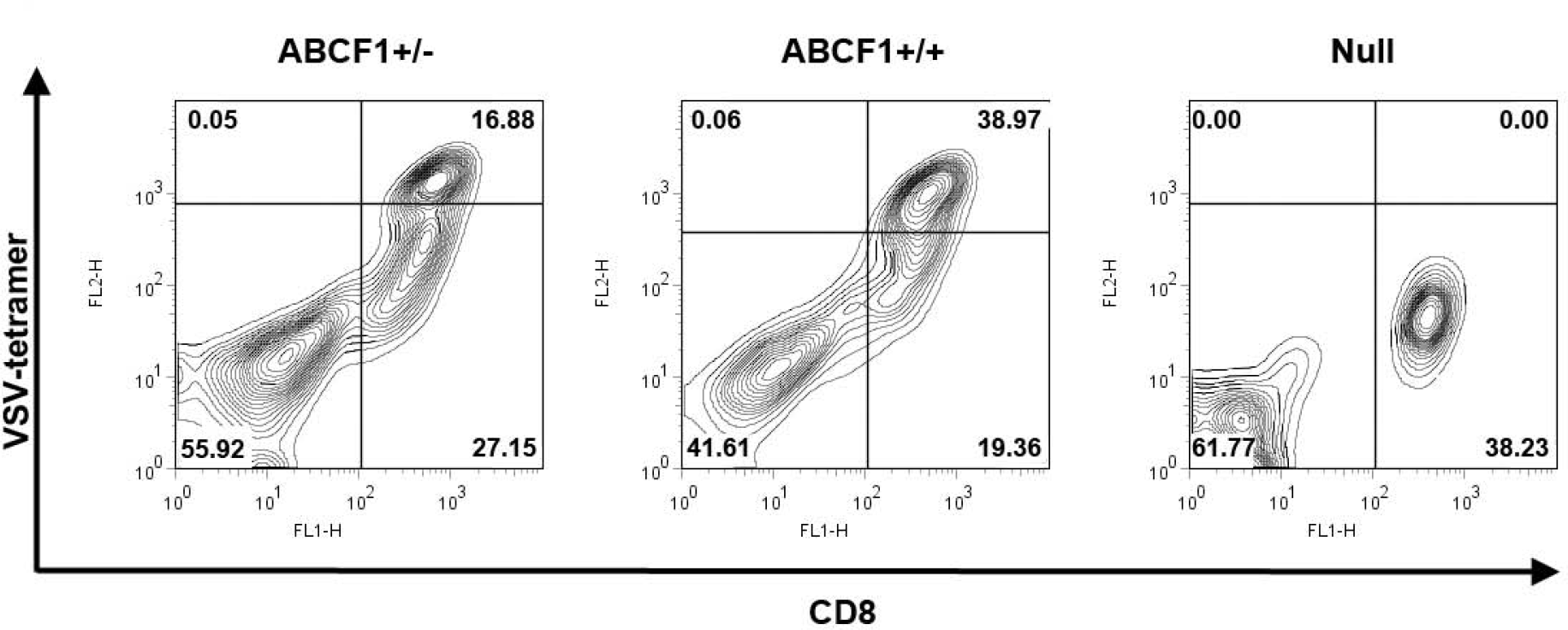
ABCF1 +/- mice produce fewer tetramer-specific CD8+ T lymphocytes after infection with VSV. Figure shown represents the results of one experiment, which was later repeated four times with similar results.

**Figure 5.**
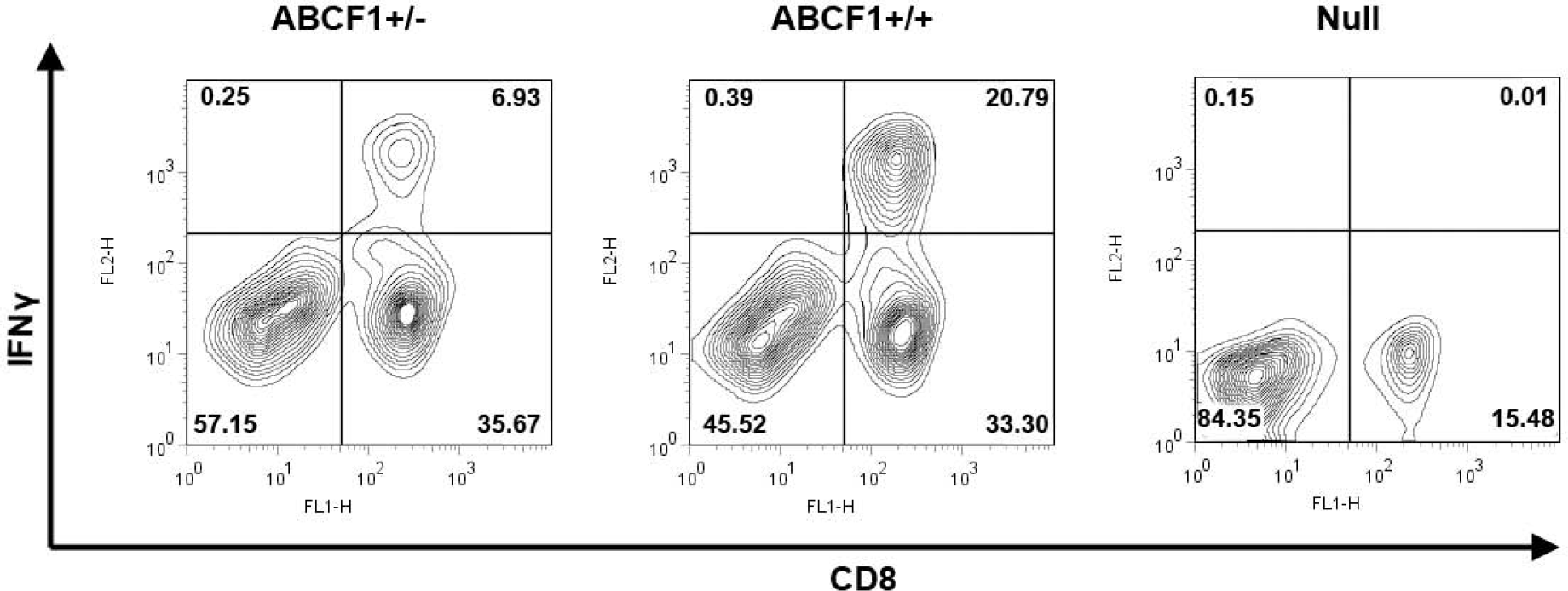
ABCF1 +/- mice produce fewer IFN**γ** positive CD8+ T lymphocytes after infection with VSV. ABCF1+/- splenocytes showed fewer IFNγ positive CD8+ T lymphocytes. Figure shown represents the results of one experiment, which was later repeated three times with similar results.

**Figure 6.**
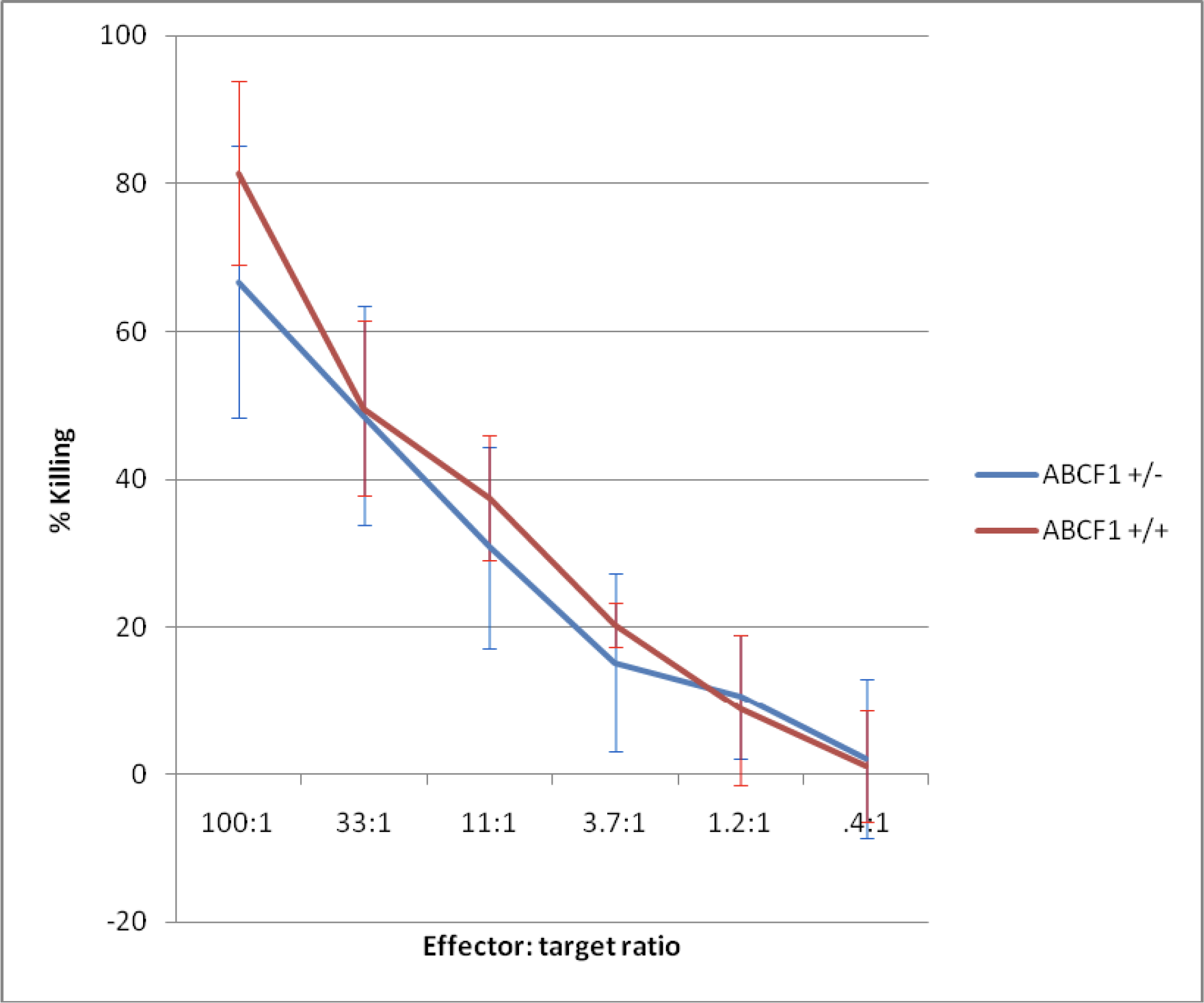
ABCF1+/- CTLs were as efficient as ABCF1 +/+ at killing of chromium. loaded VSV specific target lymphocytes. CTL activity was measured by the release of ^51^Cr into the supernatant. Figure shown represents the results of one experiment, which was later repeated three times with similar results.

To further investigate the role of ABCF1 in CD8+ cells, we used FACS to analyse their expression in the thymus and the spleen. We found that in the thymus (which contains immature and mature T lymphocytes), the population of CD8+ T lymphocytes had variable promoter activity, with quite a large percentage of cells expressing very little β-geo. This suggests that either certain subsets of CD8+cells don’t express ABCF1 or that maybe ABC is expressed upon maturation. In contrast, the spleen (which contains mature T lymphocytes), contained mostly β- geo positive cells, suggesting that ABCF1 may be involved in CD8 T lymphocyte maturation. To further test this hypothesis, FACS could be used to analyse the thymic T lymphocyte expression of fluorescein along with other markers of T lymphocyte maturation such as CD4, CD3, IL-2 receptor α, CD5, CD69 and CD44 [60]. The difference in the promoter activity in subsets of CD8+ T lymphocytes led us to ask whether ABCF1+/- CD8+ T lymphocytes could produce a functional CTL response to virus to VSV infection. ABCF1+/- mice produced fewer CD8+ T lymphocytes than their littermates. These cells produced less IFNγ than the wild type cells but were as efficient at killing target lymphocytes. Taken together, these data indicate that while ABCF1 may have a role in the development and cytokine producing ability of CD8+ T lymphocytes in response to a VSV infection. However, it does not appear to function in the CTL recognition or response to viral epitopes in the context of H-2K^b^.

Immune responses to VSV are interdependent upon TLR3 (dsRNA) and TLR7 (ssRNA) (38–41). This led us to examine the role of ABCF1 in antiviral responses triggered by polyIC that mimics dsRNA.

### ABCF1 is necessary for IFN I specific chemokine and cytokine production after Poly(I:C) stimulation

Type I IFNs are important for host defense against viruses. Once a cell is stimulated, myriad of IFN I pathway genes are produced that help in elimination of viral infection. The mechanism of their regulation and production still remains elusive. To identify if ABCF1 is necessary for this mechanism, we knocked down ABCF1 in bone marrow derived macrophages (BMDM) through RNAi and studied the levels of various IFN I pathway proteins and transcription factors. Here we activated with poly(I:C) which was used as synthetic dsRNA mimic that can activate Pattern Recognition Receptors (PRRs), including TLR3 and also RIG-I, MDA-5 and PKR.

This comprehensive approach aimed to uncover the nuanced effects of ABCF1 on cellular responses and signaling pathways.

Levels of IFN type I pathway specific cytokines, IFN-α and IFN-β were down-regulated by 2 to 3-fold when treated with *Abcf1* siRNA compared with scrambled siRNA treated group while IFN-α and IFN-β were upregulated by 3 to 4-fold in *Abcf1* over-expression group compared with scrambled siRNA (**Figured 6**). On the other hand, poly (I:C) treated group showed the highest level of IFN-α and IFN-β while *Abcf1* siRNA/ poly (I:C) group showed similar levels of cytokines in comparison with scrambled siRNA treatment group suggesting the positive effect of poly (I:C) was counteracted by the negative effect of *Abcf1* siRNA (**Figure 7**).

**Figure 7.**
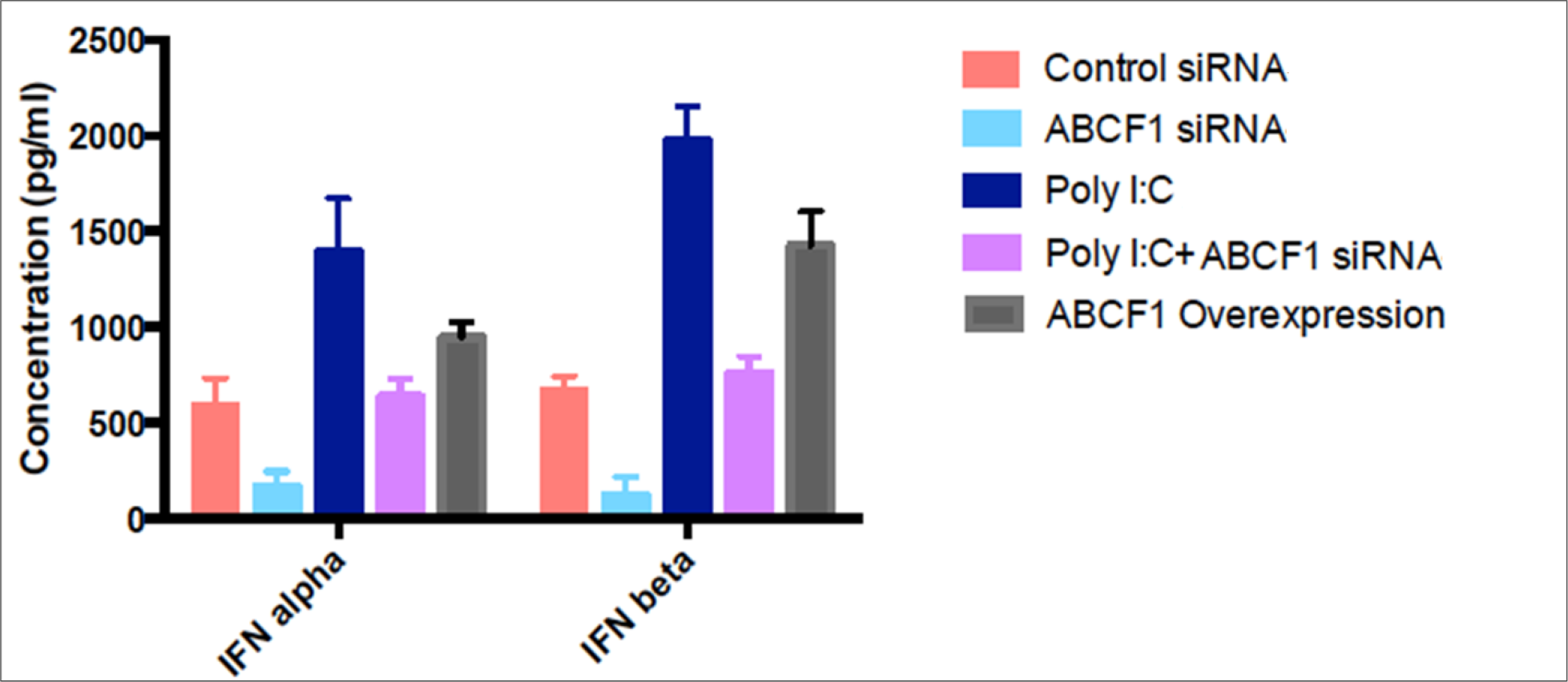
ABCF1 is essential for the production of ISG (Interferon-Stimulated Gene) specific cytokines expression of INF-**α** and **β** after Poly(I:C) stimulation. A bar graph illustrating the fold change in the levels of INF-α and β cytokines detected in the cell culture supernatants. These findings provide insights into the impact of *Abcf1*-specific siRNA treatment, both with and without Poly(I:C), on the secretion of cytokines and chemokines as well as the activation of key signaling pathways and transcription factors within BMDMs. Expression levels of the protein was assessed by western blotting analysis.

Thus, cytokine regulation by ABCF1 demonstrates that ABCF1 is likely necessary for eliminating viral infection via IFN I production. IFN-β acts as a signaling molecule that triggers the expression of OAS1 and other ISGs. OAS1, in response to the presence of viral double-stranded RNA, synthesizes 2-5A, which activates RNase L, leading to the degradation of viral RNA and inhibition of viral replication. This coordinated action demonstrates the interrelated roles of IFN-β and OAS1 in the cellular defense against viral infections (42).

### ABCF1 counteracts viral infection via JAK-STAT pathway and IRF3 production

Many signaling pathways are engaged after viral infection, leading to the formation of dsRNA (42). The cell can either inhibit translation by activating protein kinase RNA-activated (PKR) or degrade the mRNA by RNaseL by activating OAS proteins (42). Both of these pathways lead to activation and dimerization of IRF3, which lead to production of ISG’s (43, 44). To study if ABCF1 regulates this signaling pathway, phosphorylated levels of ISG proteins were analyzed in *Abcf1* siRNA treated and poly(I:C) stimulated BMDM.

BMDM treated with both *Abcf1* siRNA and poly(I:C) revealed reduced phosphorylation levels of transcription factor like NF-kB p65 (Anti NF-κB p65 antibody sourced from abcam; catalog ab16502) and IRF3 (**Figure 8A**). Reduced IRF3 dimerization was also observed in cells treated with *Abcf1* siRNA alone, whereas significant elevation in IRF3 dimerization was observed when BMDM were stimulated with poly(I:C), but levels significantly plummeted when treated with both *Abcf1* siRNA and poly(I:C) (**Figure 8B**). This suggested that ABCF1 regulates dsRNA mediated signaling via IRF3 transcription factors and thereby leads to production of ISG’s. The efficiency of *abcf*1 knocked down by RNAi by western blotting in BMDM +/- poly(I:C) was established in **Figure 7A**. This experiment and the experiments in **Figure 9** were undertaken at the same time, so the control is relevant for both figures.

**Figure 8.**
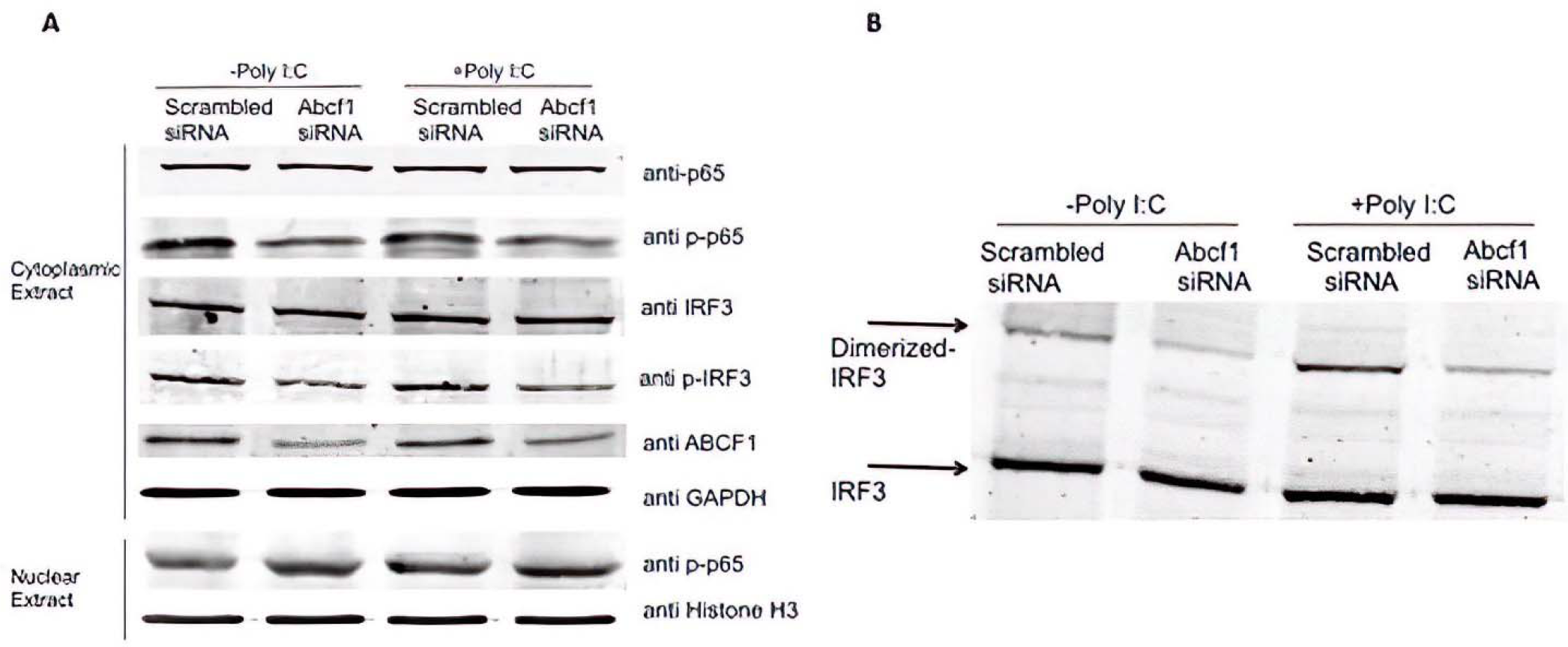
ABCF1 plays a role in regulating the phosphorylation levels of NF-**κ**B p65 and IRF3. A) Cytoplasmic and nuclear fractions obtained from BMDMs under different conditions were subjected to western blot analysis to assess the levels of transcription factors associated with Poly(I:C) stimulation confirming our hypothesis; Western Blot band sizes are as follows: anti- p65 (65 kD); anti p-p65 (65 kD); anti IRF3 (47 kD); anti p-IRF3 (36 kD); anti ABCF1 (110 kD); anti GAPDH (36 kD); anti histone H3 (15 kD); Dimerized-IRF3 (73 kD); IRF3 (36 kD) B) Whole-cell lysates were separated using native PAGE gel electrophoresis, and the analysis focused on examining the dimerization of IRF3, a transcription factor. Expression levels of the protein was assessed by western blotting analysis. Fold change calculations, as specified in the materials and methods section, were employed to quantify changes in the data. The bar graphs displayed in the figures represent the mean values along with standard deviations. The data presented here is representative of three separate experiments.

**Figure 9.**
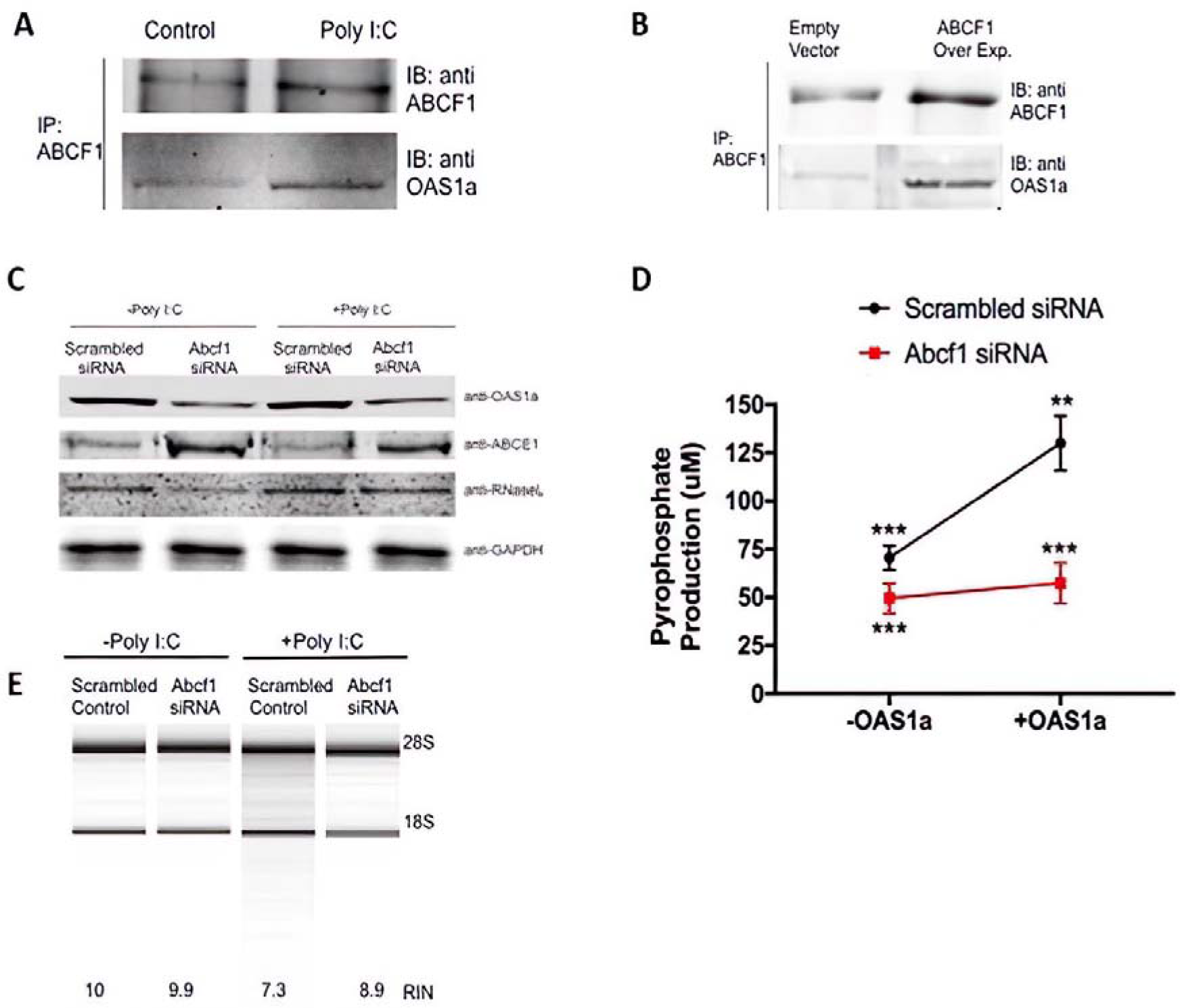
ABCF1 Parallels OAS1a Expression and Controls 2-5A Activity **A)** The immunoprecipitates were immunoblotted with anti-ABCF1 and anti-OAS1a antibodies. **B)** ABCF1 was over-expressed (Over Exp.) in BMDMs, an equal amount of whole cell lysates was immunoprecipitated with anti-ABCF1 antibody and the immunoprecipitates were detected with anti-ABCF1 and anti-OAS1a antibodies using western blots. **C)** Protein levels of OAS1a, ABCE1 and RNase L were analyzed from whole cell lysates from *Abcf1* siRNA-treated BMDMs in presence or absence of Poly(I:C). Expression levels of the protein was assessed by western blotting analysis. **D)** Increased pyrophosphate production (a readout of 2-5A activity) was measured in scrambled and *Abcf1* siRNA treated BMDMs in presence and absence of recombinant OAS1. **E)** Total RNA was extracted from *Abcf1* siRNA treated BMDMs in presence or absence of Poly(I:C) and was run on a Agilent 2100 Bioanalyzer. 28S and 18S RNA have been shown and RIN score was calculated. Bars indicate the mean ±standard deviation. The data is representative of 3 separate experiments. The expression of ABCF1 follows a pattern that parallels the expression of OAS1a, and it also exerts control over the activity of 2-5A. This suggests a regulatory relationship where ABCF1’s expression correlates with OAS1a expression and influences the activity of 2-5A.

**Figure 10.**
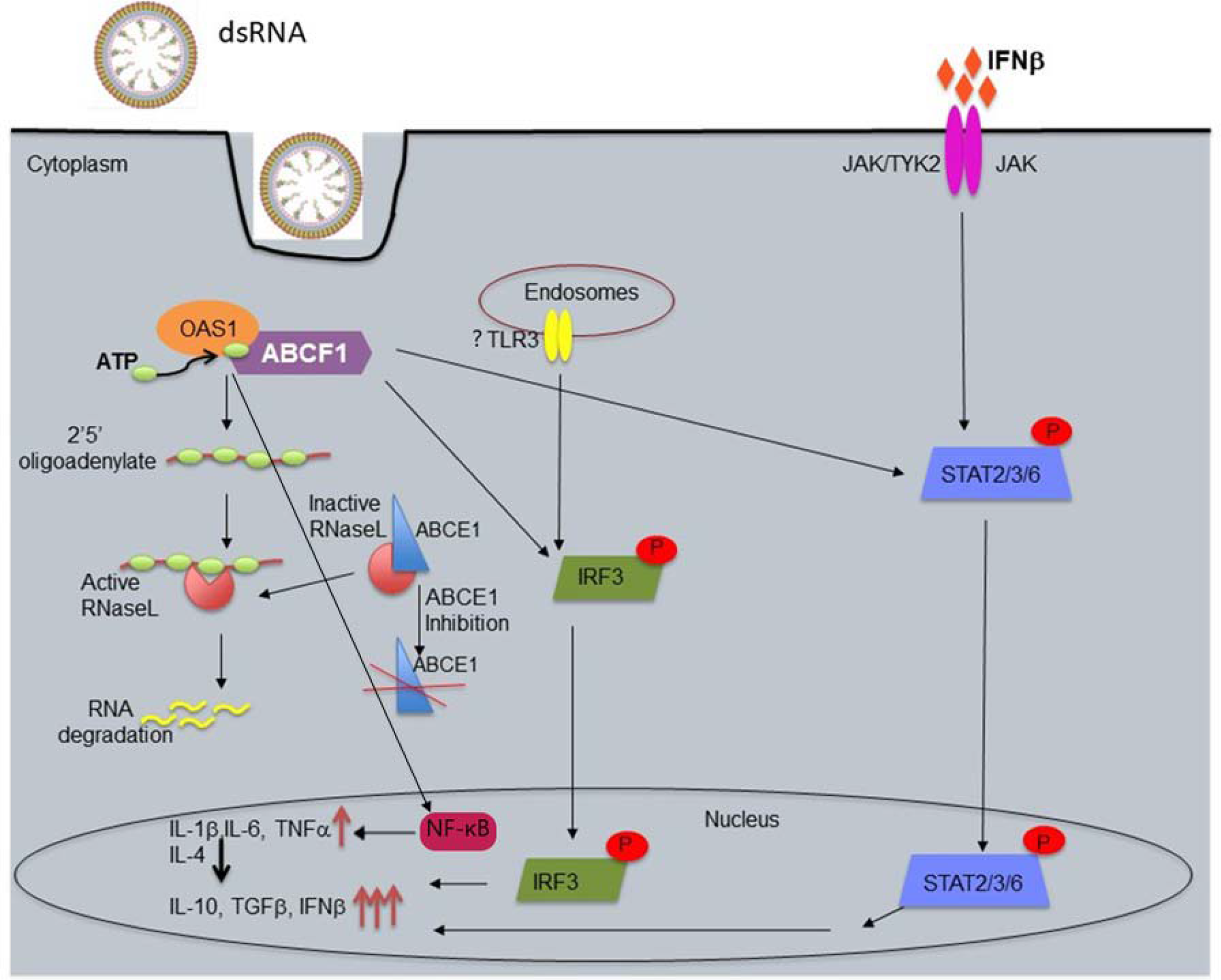
Potential Model for ABCF1 regulating dsRNA responses through a complex interplay involving OAS1a, TLR3, and JAK-STAT signaling pathways. Production of OAS1a: In response to dsRNA, one of the first antiviral proteins produced is OAS1a. OAS1a is essential for recognizing and responding to viral RNA; ABCF1 Interaction: ABCF1 associates with OAS1a upon dsRNA infection. This interaction is crucial for the subsequent steps of the antiviral response; Production of 2-5A: The binding of ABCF1 to OAS1a triggers the production of 2-5A (2’-5’-oligoadenylate), a critical antiviral molecule; Activation of RNase L: 2-5A molecules, in turn, bind and activate the enzyme RNase L. Activated RNase L degrades viral RNA, which is essential for inhibiting viral replication; TLR3 Detection: Simultaneously, dsRNA can also be detected by Toll-like receptor 3 (TLR3) located in endosomes within the cell; TLR3 Signaling: Upon activation by dsRNA, TLR3 triggers a signaling cascade that leads to the phosphorylation and dimerization of IRF3 (Interferon Regulatory Factor 3); IRF3 Nuclear Translocation: Phosphorylated and dimerized IRF3 is shuttled into the nucleus, where it functions as a transcription factor; IFN-β Production: In the nucleus, IRF3 promotes the production of interferon-beta (IFN-β) and other cytokines. IFN-β is a crucial antiviral molecule; Positive Feedback Loop: IFN-β acts in a positive feedback loop by binding to cell surface receptors and activating the JAK-STAT (Janus kinase-signal transducer and activator of transcription) pathway; ABCF1’s Role in STAT Phosphorylation: ABCF1 plays a critical role in the JAK-STAT pathway by being essential for the phosphorylation of STAT3, STAT5, and STAT6; Strong IFN-I Mediated Response: ABCF1’s involvement in STAT phosphorylation contributes to mounting a robust IFN-I (Type I interferon) mediated antiviral response, which further enhances the cell’s ability to combat the infection. This coordinated series of events demonstrates how the interplay between ABCF1, OAS1a, TLR3, IRF3, and the JAK-STAT pathway helps the host lymphocyte effectively recognize and respond to dsRNA from invading pathogens, ultimately leading to the elimination of the infection.

### ABCF1 is necessary for OAS1 and RNaseL activity after Poly(I:C) stimulation

The anti-viral OAS1-RNaseL pathway has shown to be crucial for viral RNA cleavage and production of IFN-I (45). IFN-I production initiates a feedback JAK-STAT pathway, which aids in viral elimination via phosphorylation of STAT proteins and production of ISG’s. To investigate if ABCF1 regulates OAS1a signaling, poly(I:C) stimulated BMDM were immunoprecipitated with anti ABCF1 antibody (sourced from ThermoFisher Scientific; catalog PA5-29955) and data revealed an association between ABCF1 and OAS1a (**Figure 9A, B**). BMDM were then treated with *Abcf1* siRNA in the presence and absence of poly(I:C) and OAS1a protein levels were then analyzed. Western blot analysis demonstrates a decrease in OAS1a levels when treated with *Abcf1* siRNA alone or with *Abcf1 siRNA*/poly(I:C) in comparison with the group treated with scrambled siRNA (**Figure 9C**).

To examine if ABCF1 was also necessary for OAS1 anti-viral activity, the amount of pyrophosphate (PPi) produced as a by-product of 2-5(A) formation was quantified as described in previously established assays (46–48). *Abcf1* siRNA treated and poly(I:C) stimulated BMDM showed no increase in PPi production when transfected with an expression vector encoding OAS1a or not (**Figure 9D**). On the other hand, two-fold elevation in PPi levels was seen in controls (**Figure 9D**). Total RNA was extracted from *Abcf1* siRNA treated BMDMs in presence or absence of Poly(I:C) and was run on an Agilent 2100 Bioanalyzer. 28S and 18S RNA have been shown and RIN score was calculated (**Figure 9E**).

Since downstream OAS signaling leads to RNaseL activation and viral degradation, we tested if ABCF1 regulated RNaseL levels. To investigate this, RNaseL protein levels were measured in *Abcf1* siRNA treated and poly(I:C) stimulated BMDM. Immunoblot and the regulation of RNaseL by ABCF1 is also observed without poly(I:C) stimulation (**Figure 9C**).

Previous studies have already shown that ABCE1 negatively regulates RNaseL activity (33), however, western blot analysis revealed that ABCF1 negatively regulates ABCE1 levels in BMDM treated with *Abcf1* siRNA in presence and absence of Poly(I:C) (**Figure 9C**). This confirms that ABCF1 likely positively regulates viral degradation by controlling OAS1 anti-viral activity and negatively regulating ABCE1 (**Figure 9C**).

## Discussion

The mammalian immune system has evolved to recognize and counteract many different forms of PAMPs through PRRs present either on the surface or inside the cell. This integrate mechanism is under the surveillance of many proteins, which form a vast network of signaling cascades that ultimately controls the cells reactions towards pathogenic invaders.

In our previous work we discovered that ABCF1 governs a shift in macrophage polarization. ABC transporters are a widespread group of transport proteins, constituting one of the largest and most ancient families. They are present in all currently existing biological phyla, spanning from prokaryotes to humans. However, unlike other ABC subfamilies A-D and G members, ABC subfamily F proteins lack the trans-membrane domain and do not function as transporters and are therefore, likely located in the cytoplasm and other cellular organelles. When ABCF1 was absent in macrophages, these cells exhibited impairments in internalizing TLR4 receptors and failed to switch from MyD88-dependent signaling to TRIF-dependent signaling upon LPS stimulation. Consequently, these macrophages predominantly generated pro-inflammatory cytokines specific to the NF-κB and MAPK pathways, adopting an M1-like phenotype accompanied by the M1- cytokine production. Additionally, in a sepsis model induced by endotoxins, mice haplo- insufficient for ABCF1 were unable to transition from the systemic inflammatory response syndrome (SIRS) phase to the endotoxin tolerance (ET) phase. This failure led to heightened expression of MyD88-associated pro-inflammatory cytokines in the bloodstream, activation of pyroptotic cell death pathways, and eventual renal circulatory failure, leading to premature death. These findings strongly indicate that ABCF1 expression plays a vital role physiologically. It promotes the production of TRIF-dependent anti-inflammatory cytokines and IFN-β by guiding macrophages towards an M2 phenotype, which is characterized by anti-inflammatory properties. Conversely, the absence of ABCF1 mimics MyD88 signaling, prompting macrophages to produce pro-inflammatory cytokines and adopt an M1 phenotype, which is associated with inflammatory responses (32).

Also in our previously published work, we had created a heterozygous mouse model of another ABC gene, *Abcf1*+/-, after finding the homozygous knockout mouse was embryonic lethal at 3.5 days *post coitum* (37). In a subsequent publication (32), we have demonstrated that ABCF1 is a novel E2 ubiquitin conjugating enzyme that acts as a switch between the MyD88-directed pathway and the TRIF-TIRAM directed pathway of inflammation. These pathways either lead to a pro-inflammatory cytokine secretion or anti-inflammatory cytokine secretion and control macrophage polarization from M1 to M2 phenotype. By acting as a switch between these two pathways, ABCF1 subsequently protects against sepsis-mediated cytokine storm. Interestingly, previous studies have shown the involvement of certain TRIM ubiquitin E3 ligases in LPS-TLR4 signaling (49) and regulation and manipulation of these ubiquitin E3 ligases by ubiquitin E2 conjugating enzymes (50). ATP Binding Cassette Family E member 1 (ABCE1) is a member of ATP cassette super family and is a known RNaseL inhibitor. Furthermore, studies have shown that overexpression of ABCE1 in HeLa cells inhibits IFN mediated anti-viral activity on encephalomyocarditis, thereby inhibiting 2-5A/RNaseL pathway.

In our ongoing exploration of ABCF1’s function we hypothesized that ABCF1 may play a role in viral defenses, and we assessed whether ABCF1 plays a pivotal role in CD8+ T lymphocyte that mediate anti-viral immune responses. We first examine the expression of ABCF1 within the thymus and spleen. To assess the impact of ABCF1 on lymphocyte development, flow cytometry analysis, including CD4 and CD8 staining, was conducted on the spleen and thymus (Figure 1).

The thymocyte subpopulations in ABCF1+/- showed no significant differences compared to controls. However, spleens from ABCF1+/- exhibited an elevated presence of CD4+ cells, indicative of resting CD4+ T lymphocytes, as suggested by their increased size compared to controls. Our investigation within the thymus, demonstrated a spectrum of promoter activities among CD8+ T lymphocytes (**Figure 2**). Intriguingly, a significant portion of these cells exhibited minimal β-geo expression, suggesting the interesting possibility that specific subsets of CD8+ thymocytes might either lack ABCF1 expression or activate it upon maturation (**Figure 2**).

The variation in promoter activity within specific subsets of CD8+ T lymphocytes prompted us to investigate whether CD8+ T lymphocytes with partial ABCF1 expression (ABCF1+/-) could mount an effective CTL response during VSV infection. ABCF1+/- mice yielded a reduced population of CD8+ T lymphocytes compared to their counterparts (**Figure 3, 4**). Although these cells produced lower levels of IFNγ, they exhibited equal efficiency in targeting and eliminating infected cells (**Figure 6**). In synthesizing these findings, it is evident that ABCF1 plays a pivotal role in shaping the development and cytokine-producing abilities of CD8+ T lymphocytes in response to VSV infection. However, intriguingly, ABCF1 does not appear to govern the CTL recognition or response to viral epitopes within the context of H-2K^b^ (**Figure 6**).

The immune responses triggered by VSV are intricately reliant on the cooperation between TLR3, which recognizes double-stranded RNA (dsRNA), and TLR7, which senses single-stranded RNA (ssRNA) (38–41). These fundamental interactions underpin the body’s defense mechanisms against VSV infection, Additionally, the Poly(I:C), a synthetic analog that mimics dsRNA, in the elicitation of antiviral responses has been used to understand the underlying innate immune mechanisms involved in anti-viral responses. To further unravel the complexities of these responses, we turned our attention to the role of ABCF1 in Poly(I:C)-induced immune responses. This investigation aimed to dissect the involvement of ABCF1 in the context of Poly(I:C), shedding light on its influence on the mimicry of dsRNA antiviral responses.

Poly(I:C) activates a range of PRRs, not limited to Toll-Like Receptor 3 (TLR3), but also including Retinoic Acid-Inducible Protein I (RIG-I), Melanoma Differentiation-Associated Gene 5 (MDA-5), and Protein Kinase RNA-Activated (PKR). The activation of specific PRR pathways depends on the subcellular location of poly(I:C). When present in endosomes, poly(I:C) triggers the activation of TLR3 (as demonstrated by Alexopoulou et al.(4)). In the cytosol, poly(I:C) stimulates not only RIG-I, as indicated by studies conducted by Kawai and Akira in 2008 (51) and Kato et al. in 2006 (52), but also MDA-5 (as shown by McCartney et al. (53); Kato et al. (52)), and Protein Kinase RNA-Activated (PKR, as reported by Lemaire et al. (54)).

In a study by Lee et al., (55) the response of Interleukin-6 (IL-6) to poly(I:C) was also explored. Interestingly, their findings revealed that the IL-6 response to poly(I:C) was inhibited in cells lacking RIG-I (RIG-I KO) and MDA-5 (MDA5-KO), indicating the involvement of these receptors in mediating the response to poly(I:C). Furthermore, this response was completely abolished in cells lacking the Mitochondrial Antiviral Signaling Protein (MAVS KO cells), underscoring the essential role of MAVS in orchestrating the cellular response to poly(I:C) (38–41).

In the present study, upon Poly(I:C) stimulation in BMDMs, ABCF1 is important for the production of ISG specific proteins such as IFN-β and IFN-α (**Figure 6**). Protein analysis in *Abcf1* siRNA treated BMDMs showed a decrease in NF-kB p65 phosphorylation upon Poly(I:C) stimulation, when compared with Poly(I:C) treated scrambled siRNA group (**Figure 7A**). Decreased levels of IRF3 phosphorylation and dimerization were observed in *Abcf1* siRNA treated and Poly(I:C) stimulated BMDMs (**Figure 7B**), which suggested that ABCF1 may positively regulate STAT protein activity possibly in a TRIF-dependent manner (32). Thus, cytokine-chemokine and protein phosphorylation data suggests that ABCF1 deficiency in the cells seems to phenocopy pro-inflammatory MyD88-NF-κB pathway (Phospho-NF-κB p65 antibody was sourced from abcam; catalog ab86299), whereas TRIF-IRF3 pathway was found to be ABCF1 dependent, as previously described by our lab (32).

Previous studies have shown that a cell counteracts dsRNA containing pathogens via OAS1- RNase L mediated viral degradation (3, 20, 35, 43). Our studies support the involvement of ABCF1 in regulating this signaling pathway by interacting with OAS1a and thereby controlling its 2’5’A activity (**Figure 8**). Immuno-precipitation experiments revealed that upon Poly(I:C) stimulation OAS1a was immuno-precipitated with ABCF1 (**Figure 8**). Protein analysis also showed that ABCF1 positively regulates OAS1a levels in *Abcf1* siRNA treated and Poly(I:C) stimulated BMDMs (**Figure 9**). ABCF1 was also found to negatively regulate protein levels of RNase L inhibitor, ABCE1, thus potentially positively regulating RNase L dependent RNA degradation (**Figure 9**).

The findings from our research are consistent with previous studies indicating that ABCE1 exerts a negative regulatory influence on RNaseL activity (31). Interestingly, our investigation using western blot analysis uncovered a novel facet: ABCF1 negatively modulates ABCE1 levels in BMDM subjected to *Abcf1* siRNA treatment, both in the presence and absence of Poly(I:C) stimulation (**Figure 8C**). This intriguing discovery strongly suggests that ABCF1 likely plays a positive role in enhancing viral degradation. It achieves this by not only controlling the antiviral activity of OAS1 but also by negatively regulating ABCE1, thereby fine-tuning the cellular responses against viral infections, as illustrated in **Figure 8C**. These findings underscore the intricate regulatory mechanisms orchestrated by ABCF1, shedding light on its multifaceted role in the cellular defense against viral threats.

Our data does not definitively indicate whether ABCF1 directly interacts with OAS1a or if the interaction is mediated indirectly, potentially through RNA binding. Further studies are essential to probe the nature of this interaction, clarifying whether it involves a direct physical binding or if intermediary factors, like RNA molecules, play a role. These additional investigations are crucial for a comprehensive understanding of the relationship between ABCF1 and OAS1a.

A recent study published shed light on the effect of the Neanderthal p46 OAS1 isoform on patients infected with the recent SARS-CoV-2 virus, implicated in the COVID-19 pandemic ^1^. Their study was able to demonstrate that this increased OAS1 isoform levels in patients that are in a non-infectious state are strongly associated with a reduced risk of very severe COVID-19 infection, hospitalization and susceptibility, giving the population protective immunity against the virus. The magnitude of this protective effect that was seen was particularly large, in that a 50% decrease in the odds of very severe COVID-19 infection manifestation was observed per standard deviation increase in OAS1 circulating levels. This protection was shown to be achieved through induction of OAS1 by interferons caused by the infection, which resulted in an activation of latent RNase L, which, in turn, results in a direct viral and endogenous RNA destruction (56). This was further validated by the study of Wickenhagen et al. (26), which demonstrated, using antiviral effectors encoded by IFN-stimulated genes (ISGs) screening, that OAS1 consistently inhibits SARS-CoV-2 in different contexts. Prenylation at the C-terminal is essential for OAS1 to effectively block SARS-CoV-2. This modification enables OAS1 to target intracellular areas abundant in viral dsRNA, creating a notable barrier to the virus’s replication. Moreover, this prenylation is crucial for OAS1’s ability to detect SARS-CoV-2, allowing it to selectively inhibit the virus by activating RNase L, specifically targeting the membrane organelles where the virus replicates.

Previous studies have shown that Poly(I:C) acts as an adjuvant and helps in inhibiting HIV infection. This adjuvant role elicits IFN-I production through TLR3 and shuts-down HIV replication in DCs (57). Poly(I:C) leads to the production of IFN-β, CCL2 and CXCL10 and induces a unique innate immune response against HIV. Our data suggests that ABCF1 regulates Poly(I:C) mediated IFN-I and ISG production, therefore perhaps ABCF1 could also regulate Poly(I:C) mediated TLR3 anti-viral infection after HIV exposure.

IFN-I has also been shown to induce counter-regulatory mechanisms, which has provided mechanisms to counteract chronic inflammation, viral responses and even cancer (58). IFN-I regulates these inhibitory factors by modulating Programmed Death-Ligand 1 (PD-L1) and Indoleamine 2,3-dioxygenase (IDO) levels during Lymphocytic Choriomeningitis Virus (LCMV) infection ^51^ and counteracts metastases and T lymphocyte inhibitory signals and promotes T lymphocyte responses in cancer. In melanoma and breast cancer, IFN-I has been known to drive the expression of TNF-related Apoptosis-Inducing Ligand (TRAIL), which triggers caspase 8 mediated cellular apoptosis (59, 60). As ABCF1 regulates IFN-I and it is a target of caspase 8 (61), perhaps ABCF1 both regulates the anti-viral response against LCMV and acts to restrict cancer metastases.

Our data defines a mechanism for control of OAS1, that has recently demonstrated to play a crucial protective role in SARS CoV-2 infections. We have identified a mechanism that regulates innate immune signaling. It operates independently of its E2 Ubiquitin-conjugating activity and does so by docking with and interacting with OAS1. This interaction mediates the phosphorylation of OAS1, which then potentially initiates its anti-viral activity through modulating RNaseL activity via ABCE1. Furthermore, we find that ABCF1 regulates the cytokine production via IRF3 phosphorylation and dimerization which modulates anti-viral JAK-STAT signaling via a positive feedback loop through IFN-β and phosphorylation of STAT transcription factors that enhance viral clearance. Overall, these studies define ABCE1 and/or ABCF1 as drug targetable proteins to increase phosphorylation of OAS1 that could act as anti- viral agents for SARS CoV-2 and other viruses such as VSV.

## Materials and Methods

### Mice and cells

*Abcf1*^+/−^ mice (Het; B6.Cg-Abcf1 < Gt(XK097)Byg) mice were generated as previously described (37) and C57BL/6J (Jackson Laboratories; 000664) mice were used as wild-type (WT) controls for *in vivo* experiments. All *in vivo* experiments were approved and performed in accordance with The University of British Columbia Animal Care Committee and Canadian Council for Animal Care (CCAC).

### Response to Poly I:C after ABCF1 siRNA knockdown or overexpression in Bone Marrow-Derived Macrophages

Bone Marrow-Derived Macrophages (BMDM) were extracted from the femur and tibia of C57BL/6 mice and then differentiated into macrophages according to the methodology outlined in reference (62). To summarize, bone marrow was obtained from C57BL/6 mice by flushing their tibia and femur with Roswell Park Memorial Institute medium (RPMI) enriched with 10% heat-inactivated Fetal Bovine Serum (FBS). These bone marrow cells were cultured in 10 ml of RPMI supplemented with 10% FBS, glutamine, and 30% L929 cell supernatant, containing macrophage colony-stimulating factor. The initial cell density was set at 1x10^6^ cells/ml, and the cultures were maintained in 100-mm Petri dishes at 37°C with 5% CO2 in a humidified environment. Gene knockdowns in BMDM cells were achieved through RNA interference (RNAi) using Lipofectamine™ RNAiMAX Transfection Reagent from ThermoFisher Scientific (catalog number 13778075), following the manufacturer’s guidelines. Approximately 2x10^7^ BMDM cells were transfected with RNAiMAX and 375 pmoles of *Abcf1* or scrambled siRNA, incubated for 48 hrs, and subsequently treated with 10 μg/ml of PolyI:C for 24 hrs. The efficacy of the knockdown was assessed through Western blot analysis. Specifically, knockdown of the *Abcf1* gene utilized Mouse *Abcf1* siRNA from Santa Cruz Biotechnology (catalog number sc-140760), resulting in a consistent reduction of 79-81% compared to control samples treated with scrambled siRNA. Conversely, over-expression studies were conducted using a mouse *Abcf1* construct expressed in the pcDNA 3.1 vector/Lipofectamine™ 2000 Transfection Reagent from ThermoFisher Scientific (catalog number 11668027), following the manufacturer’s instructions. Approximately 2x10^7^ cells were transfected with Lipofectamine 2000 and 30 μg of *Abcf1* overexpression vector, incubated for 48 hrs, and subsequently treated with 10 μg/ml of PolyI:C for 24 hrs. The success of the overexpression was again verified through Western blot analysis.

ABCF1 expression was manipulated in this study by employing two distinct approaches. Firstly, ABCF1 was overexpressed (designated as Over Exp.), where the cellular levels of ABCF1 were intentionally increased. Secondly, specific small interfering RNA (siRNA) was utilized to downregulate ABCF1 expression in Bone Marrow-Derived Macrophages (BMDM) for 48 hours. After treatment with siRNA or ABCF1 overexpression for 48 hours, the BMDM were incubated with Poly I:C, a synthetic analog of double-stranded RNA, at a concentration of 10μg/ml. This treatment was applied for a duration of 24 hours, allowing for the observation of cellular responses under the influence of Poly I:C (sourced from Invivogen, San Diego, CA, USA). The knockdown of the *Abcf1* gene utilized mouse *Abcf1* siRNA, resulting in a consistent reduction of 79-81% in protein expression compared to control samples treated with scrambled siRNA as assessed by western blotting. ABCF1 was overexpressed was also assessed by western blotting. In essence, the study explored the impact of ABCF1 manipulation under varied conditions, encompassing both overexpression and siRNA-mediated knockdown, in the presence and absence of Poly I:C stimulation.

### Vesicular stomatitis virus studies

The vesicular stomatitis virus (VSV), Indiana Strain, was given as a gift from Frank Tufaro (University of British Columbia, Vancouver, Canada). VSV was cultured on Vero cells (ATCC) and the titre was determined by plaque assays and reported as Tissue Culture Infectious Dose affecting 50% of the culture (TCID)50 [11]. The CTLs were generated by inoculating ABCF1+/- mice and their littermate controls with 1x10^5^ TCID50 of VSV. Their spleens were removed seven days later and mashed through a 40 μm cell strainer produce a single cell suspension. Splenocytes were then washed two times with PBS and incubated for 5 days at 37°C in 25 mL CTL medium (RPMI-1640 containing 20 mM HEPES, 1 mM Na-pyruvate, 0.1 mM non-essential amino acids, 100 IU/mL penicillin, 100 μg/mL streptomycin and 10% heat-inactivated FBS (Hyclone)) per spleen (approximately 10^6^ cells/mL) with 1 μM of VSV-NP52-59 peptide (RGYVYQGL) [12]. On day 5, the CTLs were harvested, washed three times, counted, resuspended (using the initial undiluted concentration of 5 x 10^6^ cells/mL) and mixed with RMA-S target T lymphocytes at the indicated ratios. RMA-S is a thymoma cell line that presents the VSVNP52-59 peptide in the context of H-2K^b^ [13]. RMA-S were counted and resuspended at 2 x 10^6^ cells/mL in CTL media for each sample. The RMA-S were then either pulsed by the addition of 1 μL with VSV-NP52-59 peptide or left un-pulsed as a negative control. 10 μL (100 μCi) of ^51^Cr (sodium chromate, Amersham Biosciences) was added to each sample and incubated at 37°C for 1 hr. The cells were then washed three times in PBS, counted and mixed with the effector CTLs at the indicated ratios for 4 hr at 37°C. For maximal and minimal counts, RMA-S cells were either lysed with 5% Triton X-100 or incubated in media lacking the effector cells. Following the incubation, the cells were pelleted and 100 μL of supernatant was removed. The released 51Cr was then quantified in a gamma counter (LKB Instruments, Gaithersburg, MD). The specific killing was quantified using the formula: ((experimental - minimum control) / (maximum – minimum control)) x 100%. 100 μL of splenocytes from the expanded culture were plated per well into a 96 well U bottom plate and washed three times with FACS buffer, The cells were then stained for their expression of CD8, CD4, IFN-γ and H-2K^b^-NP specific T lymphocyte receptors (using Phycoerythrin (PE) labelled-H-2K^b^-NP tetramers ((immunomics-BeckmanCoulerTM)). For Fluorescein isothiocyanate (FITC)-αCD8 (Pharmingen, Oakville, ON) and PE-αCD4 (Pharmingen, Oakville, ON) staining, a 1:100 ratio of antibody to FACS buffer was used. For PE labelled-H-2K^b^-NP tetramer staining, a 1:10 ratio of tetramer to FACS buffer was used. Cells were incubated with antibodies at 4°C for 20 min on ice. For IFN-γ staining, the original 100 μL of expanded splenocytes were initially incubated in the presence of GolgiStop (Pharmingen, Oakville, ON), and then washed twice with FACS buffer. The cells were then fixed by incubating them in fix buffer (4% paraformaldehyde) for 20 min on ice. Once fixed, the cells were stained as described above with FITC-αCD8. Cells were then permeabilized using a permeabilization buffer containing 0.1% saponin, 1% fetal calf serum, 0.1% sodium azide in PBS. PE-αIFN-γ antibodies (Pharmingen, Oakville, ON) were diluted 1:100 in FACS buffer, added to the cells in permeabilization buffer and incubated at room temperature for 20 min. Finally, the cells were washed once in permeabilization buffer and resuspended in fix buffer. All of the stained cells were analyzed on the FACS scan (described above).

### Transfection studies

Gene knockdowns in cells were achieved through RNA interference (RNAi) using Lipofectamine™ RNAiMAX Transfection Reagent from ThermoFisher Scientific (catalog number 13778075), following the manufacturer’s guidelines. SiRNAs pools were generated by Santa Cruz Biotechnology are a proprietary pool of 3 target-specific 19-25 nucleotides siRNAs. The effectiveness of the knockdown was assessed through Western blot analysis. Specifically, knockdown of the ABCF1 gene was accomplished using Mouse *Abcf1* siRNA obtained from Santa Cruz Biotechnology (catalog number sc-140760), consistently resulting in a reduction of 79-81% compared to control samples treated with scrambled siRNA. For other genes, including mouse cIAP2 and mouse TRIF, siRNAs from Santa Cruz Biotechnology (catalog numbers sc- 29851 and sc-106845, respectively) were used to achieve knockdown. Complementary, gene over-expression studies were carried out using a mouse abcf1 construct expressed in the pcDNA 3.1 vector was purchased from GenScript (Clone Id: Omu73202; XM_006524122.2).

Lipofectamine™ 2000 Transfection Reagent from ThermoFisher Scientific (catalog number 11668027), following the manufacturer’s instructions. The success of the overexpression was verified by conducting Western blot analysis in transfected cells were used as the control.

#### ELISA Cytokine Analysis

To analyze cytokine levels, we prepared cell culture supernatant from BMDM 2X10^7^ cells. Treatments and incubations were conducted as specified in the figure legends of the respective experiments, and the cytokine analysis was carried out in accordance with the manufacturer’s instructions for mouse R and D Systems for INF-α and β (Catalog #: 42120-1 and Catalog #: 42400-1). The resulting cytokine levels were quantified and visually represented using bar graphs.

#### Co-immunoprecipitation

A total of 2X10^7^ cells were subjected to lysis using a freshly prepared lysis buffer containing the following components: 20 mM Tris HCl at pH 8.0, 137 mM NaCl, 1% Nonidet P-40, 2 mM EDTA, 1X Halt protease and phosphatase inhibitor cocktail obtained from ThermoFisher Scientific (catalog number 78440), and 20 mM N-Ethylmaleimide sourced from Santa Cruz Biotechnology (catalog number sc-202719). 2ug of specific antibody was incubated for 1 hour and then Protein A or Protein G beads were utilized and initially washed twice with a wash buffer containing 10 mM Tris HCl at pH 7.4, 150 mM NaCl, 1% Nonidet P-40, 1 mM EDTA, 1X Halt protease and phosphatase inhibitor cocktail, and 20 mM N-Ethylmaleimide. Subsequently, these beads were centrifuged at 3,000xg for 2 minutes at 4°C and then incubated with the specified antibody for a duration of 4 hours at 4°C while being gently rotated on a shaker. Following the antibody incubation, the beads were washed twice and then incubated with the cell lysate overnight at 4°C. After this incubation, the beads were washed again, and the protein complex was eluted by acidification. This was achieved by applying 3 x 50 μl of 0.1 M glycine at pH 2 to the sample, followed by incubation for 10 minutes with frequent agitation before gentle centrifugation. Finally, the eluate was neutralized by adding an equal volume of Tris HCl at pH 8.0.

#### Western blotting

Cell lysates were prepared using the method previously described. Briefly, the cells were lysed in RIPA buffer (1xTris buffered saline, Nonidet P40, 0.5% sodium deoxycholate, 0.1 sodium dodecyl sulphate (SDS), 0.004% sodium azide, Santa Cruz Biotechnologies) with HALT protease and phosphatase inhibitor cocktails (Thermo Scientific) on ice for 40 minutes with vortexing every ten minutes. Subsequently cells were centrifuged at 15,000 x RCF for 5 minutes and supernatant was collected. Total protein was quantified using a Bradford assay and measured using the Molecular Devices Vmax kinetic micro plate reader. A total of 50μg of protein, in 20μL of 1x NuPAGE SDS sample buffer (Thermo Scientific) and was heated to 95 ° for 5 minutes, before being separated by SDS polyacrylamide electrophoresis (PAGE). Resolved samples were transferred to nitrocellulose membrane (Bio-Rad) before being blocked in 5% (w/v) skim milk with 0.2% Tween 20 (Bio-Rad). Loading was additionally controlled by detecting GAPDH as a control (Anti GAPDH antibody sourced from abcam; catalog ab181602).

For all co-immunoprecipitation (Co-IP) samples, the sample buffer was formulated without dithiothreitol (DTT), unless explicitly stated otherwise (Anti ABCF1 antibody for co-immunoprecipitation was sourced from Proteintech; catalog 13950-1-AP). In contrast, all other samples intended for Western blot analysis were prepared with DTT included in the sample buffer. The lysates were subjected to electrophoresis and subsequent immunoblotting using 2ug of the specified antibodies. Western blot experiments were conducted in triplicate to ensure consistency and reliability.

#### Pyrophosphate measurements

Pyrophosphate (Ppi) produced as a by-product of 2-5(A) formation was quantified as described in previously established assays (44–46).

#### IRF3 dimerization assay

Anti immune r-IRF3 antibody was sourced from Cell Signaling Technology (catalog 4947)A total of 2X10^7^ cells were subjected to lysis using a buffer composed of 50 mM Tris at pH 8.0, 1% NP40 (abcam; ab142227), 150 mM NaCl, 1 mM PMSF (Santa Cruz Biotech; sc-482875), and 20 μg/ml of aprotinin. Additionally, the lysates were enriched with native PAGE sample buffer containing 125 mM Tris at pH 6.8 and 30% glycerol. Following this, the samples were separated using native PAGE and subsequently subjected to analysis via immunoblotting.

#### Cytoplasmic and Nuclear fractionation

Macrophages derived from bone marrow were exposed to the specified treatments. Subsequently, the separation of cytoplasmic and nuclear fractions was carried out using the NE-PER Nuclear and Cytoplasmic Extraction Reagents from Thermo Scientific (catalog number 78833), following the guidelines provided by the manufacturer.

#### Bar graph generation and Fold change calculation

Bar graphs were generated using GraphPad Prism software from San Diego, CA. To calculate the fold change of cytokine and 29mmune29r-kinase levels in *Abcf1* siRNA-treated BMDMs (with or without PAMP treatment), we normalized the mean pixel density to scrambled siRNA-treated BMDMs (with or without PAMP treatment), respectively. Similarly, to determine the fold change of cytokine and phospho-kinase levels in scrambled siRNA- and PAMP-treated BMDMs, we normalized the mean pixel density to scrambled siRNA- and non-PAMP-treated BMDMs. This approach allowed us to quantify and compare the changes in cytokine and phospho-kinase levels under different experimental conditions.

#### Statistical Analysis

All experiments were repeated three times and the means and statistical significance determined. Normally distributed Western blot data was subjected to the appropriate t test (paired) statistical test (using Graphpad prism software, version 10.1.2(324) with 95% confidence intervals. p values less than 0.05 were considered significant. All data were represented as mean ± SEM.The data are expressed as the mean ± standard deviation (SD). The sample size for each experiment is specified in the figure legend, indicating the number of observations or data points used in the analysis.

## Acknowledgements

We express our gratitude to Drs. Giorgia Caspani and Eliana Al Haddad for their valuable comments on the manuscript. The research reported in this work was financially supported by operating grants awarded to WAJ from the Canadian Institutes of Health Research (CIHR), specifically grants MOP-86739 and MOP-133634 and by donations to the laboratory of WAJ through the Sullivan Urology Foundation at Vancouver General Hospital (https://www.urologyfoundation.ca). Additionally, we acknowledge support through a Collaborative Research Agreement (CRA) with Mynd Life Sciences, which was instrumental in advancing our research at the University of British Columbia. Furthermore, H.A. received support in the form of a studentship from the Centre for Blood Research.

## Author Contributions

Conceived and Supervised Project: WAJ Designed research: SW, HA, CGP, WAJ Performed research: SW, HA, LM, KC Analyzed data: SW, HA, LM, CGP, KC, SK, WAJ Wrote paper: SW, HA, WAJ Edited paper: CGP, KC, SK, WAJ

## Materials and Correspondence

For any correspondence or requests related to this research, please direct them to WAJ.

