## Supplementary data for "An ATP-Binding Cassette Transporter Gene Links Innate and Adaptive Immune Responses"

**Appendix**

### Statistics summary

- Scrambled siRNA and Abcf1 siRNA groups (with or without LPS stimulation) were compared using paired t-test. A p value of 0.05 or below is considered to be statistically significant.
- Without any LPS stimulation, abcf1 siRNA downregulated abcf1 by 69.2% and this downregulation was significantly different than scrambled siRNA treatment with p value of 0.0242
- With LPS stimulation, abcf1 siRNA downregulated abcf1 by 50.2% and this downregulation was significantly different than scrambled siRNA treatment with p value of 0.0186

Supplementary Figure 1

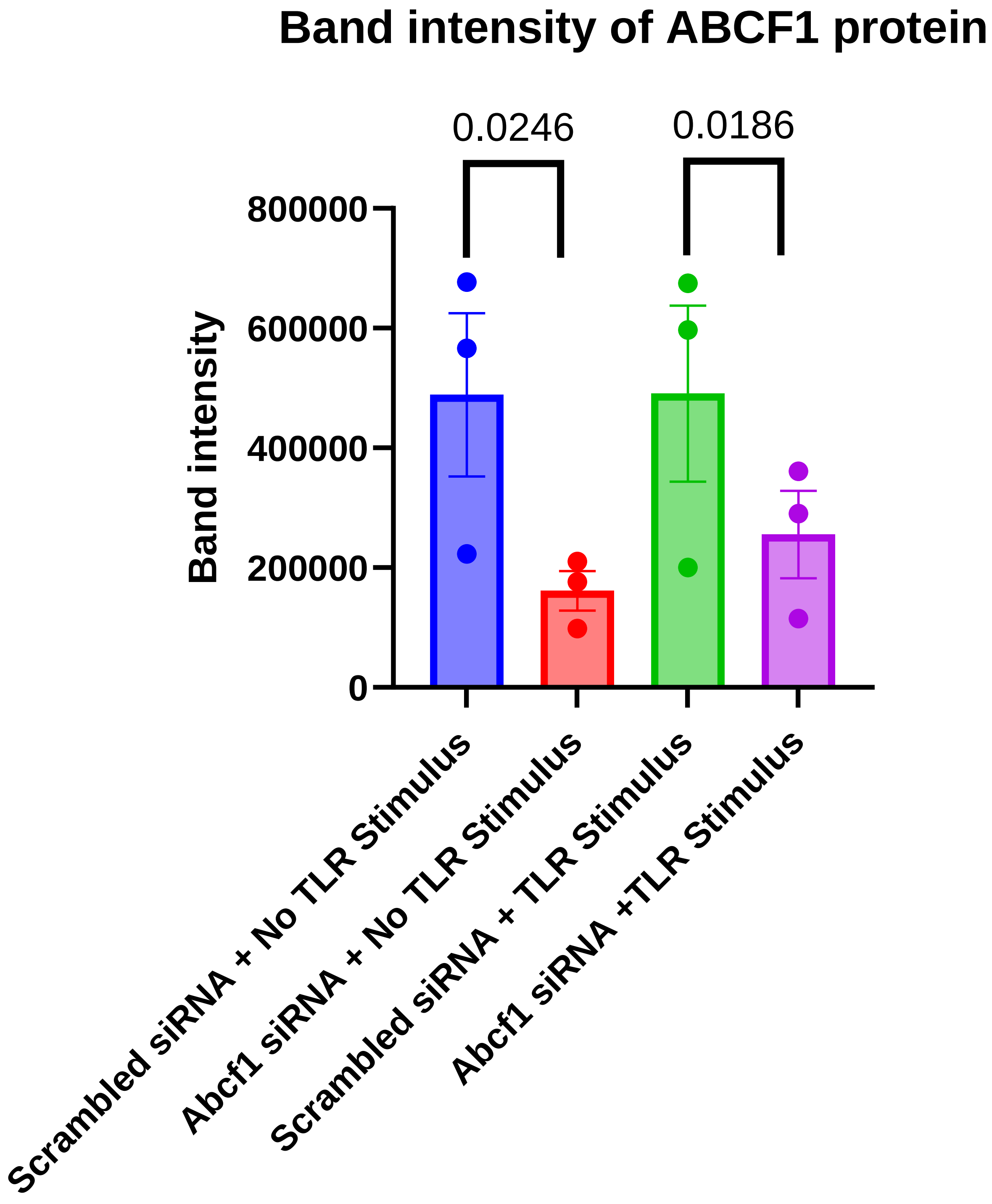


Supplementary Figure 2

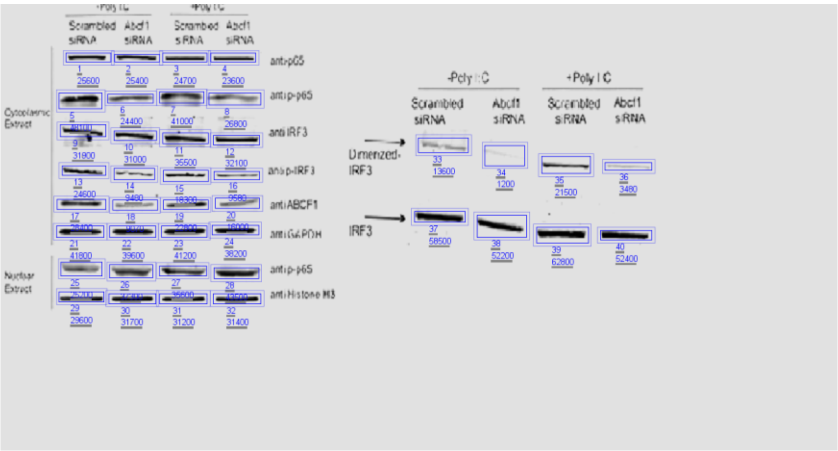


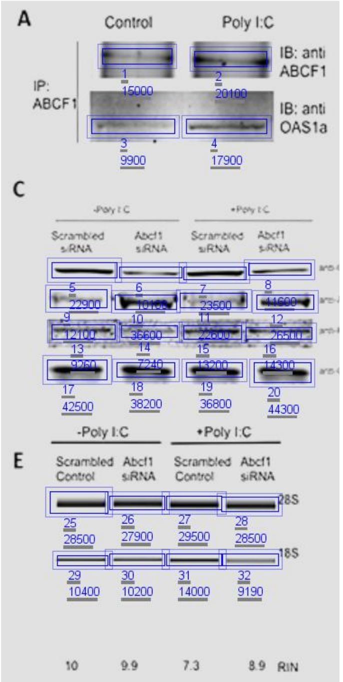


Numbers of density for the protein bands for fig7 and fig8

Supplementary Figure 3
